# Unravelling historical, taxonomic, and cultural influences on the etymology of scientific names across Animalia

**DOI:** 10.64898/2026.01.31.703012

**Authors:** Kota Nojiri, Keito Inoshita, Haruto Sugeno, Takumi Taga

**Affiliations:** Graduate School of Agricultural and Life Sciences, The University of Tokyo, 1-1-1 Yayoi, Bunkyo, Tokyo 113-8657, Japan; The University Museum, The University of Tokyo, 7-3-1 Hongo, Bunkyo, Tokyo 113-0033, Japan; Faculty of Business and Commerce, Kansai University, 3-3-35 Yamate-cho, Suita, Osaka, Japan; Data Science and AI Innovation Research Promotion Centre, Shiga University, 1-1-1 Baba, Hikone, Shiga, Japan; Faculty of Symbiotic Systems Science, Fukushima University, 1 Kanayagawa, Fukushima, Fukushima, Japan; Graduate School of Pharmaceutical Sciences, Nagoya University, Furo-Cho, Chikusa, Nagoya, Japan

**Keywords:** epithets, etymology, animalia, scientific name

## Abstract

Animal naming is fundamental to scientific communication, yet it also reflects the historical and cultural contexts in which names are bestowed. Scientific names function as taxonomic labels and enduring records of human engagement with nature. Owing to this dual role, species names have recently attracted increasing attention from historical and humanities perspectives, both for their informative value and for the biases they may encode. To objectively assess these patterns at a large scale, we investigated etymological trends across Animalia using a comprehensive dataset of species names. Our analyses reveal that naming practices are shaped by a combination of historical events, taxonomic traditions, and cultural influences. Major global disturbances coincided with marked declines in species descriptions, whereas advances in biological techniques were associated with shifts in naming practices. Furthermore, etymological trends differed among phyla, indicating that taxonomic communities vary in their naming conventions. These differences suggest that taxonomists’ preferences, shared aesthetics, available knowledge, and cultural biases are differentially preserved in scientific names. Together, our results demonstrate that zoological nomenclature constitutes a valuable archive for understanding the historical and cultural dimensions of taxonomy.

## 1. Introduction

The classification of natural entities is one of the most fundamental scientific practices, providing the conceptual foundations through which humans make sense of nature.

Among the many systems devised to recognise and organise objects, such as the periodic table of the elements and stellar nomenclature, the Linnaean system known as the binomial nomenclature has provided a stable and universal framework for defining scientific names of organisms for over two and a half centuries. For animals, the International Code of Zoological Nomenclature (hereinafter, the Code) is widely accepted as the governing framework for the binomial nomenclature [1].

The epithets (specific names) that comprise scientific names encapsulate a wide range of information, including morphological traits, ecological and behavioural features, geographic origins, cultural references, and the names of people associated with the research or the organism [2–10]. In this sense, scientific names serve not only as taxonomic labels but also as records of how humans have observed, interpreted, and valued organisms. Far from being purely descriptive, species epithets reflect the aesthetics, knowledge, and biases of the individual scientists and scientific communities who bestowed them. As such, naming practices constitute a cultural and historical archive of human engagement with nature, embedded within the formal structure of taxonomy [11].

Recently, several aspects of this archive have become subjects of academic and public attention in zoological nomenclature. For instance, gender imbalance in eponyms (names honouring specific persons) [7,10,12] has highlighted systematic disparities in who is commemorated through taxonomic practice. Similarly, epithets referencing historically controversial individuals have prompted broader societal debates concerning the values embedded in scientific naming. More broadly, such discussions align with arguments in the history and sociology of science that scientific practices are not value-neutral but are shaped by social, cultural, and institutional contexts. Londa Schiebinger has argued in the history of science that systems of classification and knowledge production often encode implicit biases, reflecting the perspectives and power structures of the communities in which they are developed [13]. From this perspective, controversies surrounding eponyms can be understood as part of a wider pattern in which taxonomic names function not only as scientific labels but also as cultural artefacts. These discussions have led to calls for revision of the Code [12,14–17] or for renaming species whose etymology is considered ethically problematic [14,18].

However, the foundational principle in zoological nomenclature defined by the Code is the preservation of scientific stability, and taxonomic communities generally oppose renaming except under exceptional circumstances [1]. Consequently, many scientists have objected to these criticisms [19–24]. We refer to these debates only to emphasise that historically taken for granted naming practices are increasingly viewed as worthy of quantitative scrutiny. Understanding long-term naming trends provides essential context for current debates on taxonomic culture and for assessing changes in naming practices over time [25].

Despite the conceptual and social importance of naming, systematic studies of species-name etymology remain sparse. Previous studies have focused primarily on specific clades, such as helminths [10], phytophagous arthropod groups [4], molluscs [7], planarians [2], rotifers [6], and spiders [8]. These studies revealed notable patterns: a historical decline in morphology-based names, an increase in geography- and people-based epithets [3,5,8,26]. Moreover, they have indicated that cultural factors influence naming decisions and underscored that nomenclature is shaped by far more than biological traits alone [8]. Yet, it remains unclear whether the trends reported in taxon-specific studies reflect localised phenomena or general patterns across Animalia. Addressing this question requires a comparative analysis that spans the major phyla of animals.

However, manual etymological classification is extremely labour-intensive, requiring the examination of original descriptions for a vast number of species [8]. Consequently, even the largest existing etymological datasets remain taxonomically restricted, and reconstructing naming trends at the scale of Animalia has long been impractical.

Recent methodological advances, however, have begun to overcome this barrier. Several studies demonstrated that large language models (LLMs) can categorise species epithets into pre-set etymological labels with accuracies of around 90%, establishing the feasibility of large-scale, semi-automated etymological annotation [3,5,26]. Their results show that LLMs can reliably distinguish major etymology types such as morphology, geography, and people. Importantly, these models allow for the first quantitative reconstruction of naming practices across the full breadth of Animalia, a task unattainable through human effort alone. This methodological innovation opens the door to understanding naming not merely as a technicality of taxonomy but as an evolving cultural practice embedded in scientific history.

Here, we present the first Animalia-wide analysis of species-name etymology. Using a large-scale dataset annotated by an LLM, we quantify the historical dynamics of six etymological categories of naming over more than 250 years of taxonomic history. We further compare these patterns across major animal phyla to evaluate the naming culture in different taxa. In addition to examining temporal trends, we incorporate information on the geographic origins of authors to evaluate how the diversification of the global taxonomist community has influenced naming practices. As taxonomic authority gradually expanded beyond its historically Eurocentric core, scientists from Asia, South America, Africa, and other regions may begin to introduce new naming preferences.

Our goal is not to assess the appropriateness of particular names, nor to make normative claims about how species should be named. Rather, we aim to provide a rigorous empirical foundation for understanding how humans have conceptualised, categorised, and commemorated the animals they describe. Through this integration of etymological, historical, and author-origin data, our study offers a new perspective on how global shifts in scientific participation have shaped the cultural evolution of zoological nomenclature.

## 2. Methods

### (a) Dataset

We first obtained an available dataset of epithets compiled by Mammola et al., which includes 48,464 epithets of spiders and etymological labels [8]. We then acquired a comprehensive list of all valid animal species names from the open-source dataset, Catalogue of Life (CoL) [27]. This dataset includes information on phylum, genus (generic names), epithets, author, and year of publication. Only currently accepted names were used, and records lacking year information were removed prior to analyses.

### (b) Establishing an etymology category framework

Previous studies on etymological analyses of scientific names have typically relied on categories, often based on subjective interpretation [4–6,8,9,26]. As a result, category definitions and boundaries have varied across studies, limiting direct comparability among datasets. It is therefore necessary to establish a quantitatively grounded and internally consistent classification framework before analysing Animalia-wide naming patterns.

We conducted a preliminary embedding-based semantic clustering using the spider dataset to evaluate the etymology labels traditionally used in previous studies: *Morphology*, *Ecology & Behaviour*, *Geography*, *People*, *Culture*, and *Other*. For each epithet, a 3000-dimensional semantic vector was generated using the GPT-4o-mini model. K-means clustering was performed, identifying seven coherent semantic clusters. We selected the 50 epithets closest to the centre of each cluster as representative examples to interpret the semantic meaning of each cluster. For visual interpretation, t-distributed stochastic neighbour embedding (t-SNE) was applied to a low-dimensional representation obtained by independent component analysis (ICA), which revealed clear substructure within *Morphology*, whereas no distinct clustering was observed for *Ecology & Behaviour* or *Culture* (Figure S1). Based on these results, we established a refined set of etymological labels for epithets: *Abstract Morphology*, *Specific Morphology*, *Conceptual Morphology*, *Geography*, *People* and *Other* (Table 1).

**Table 1.**
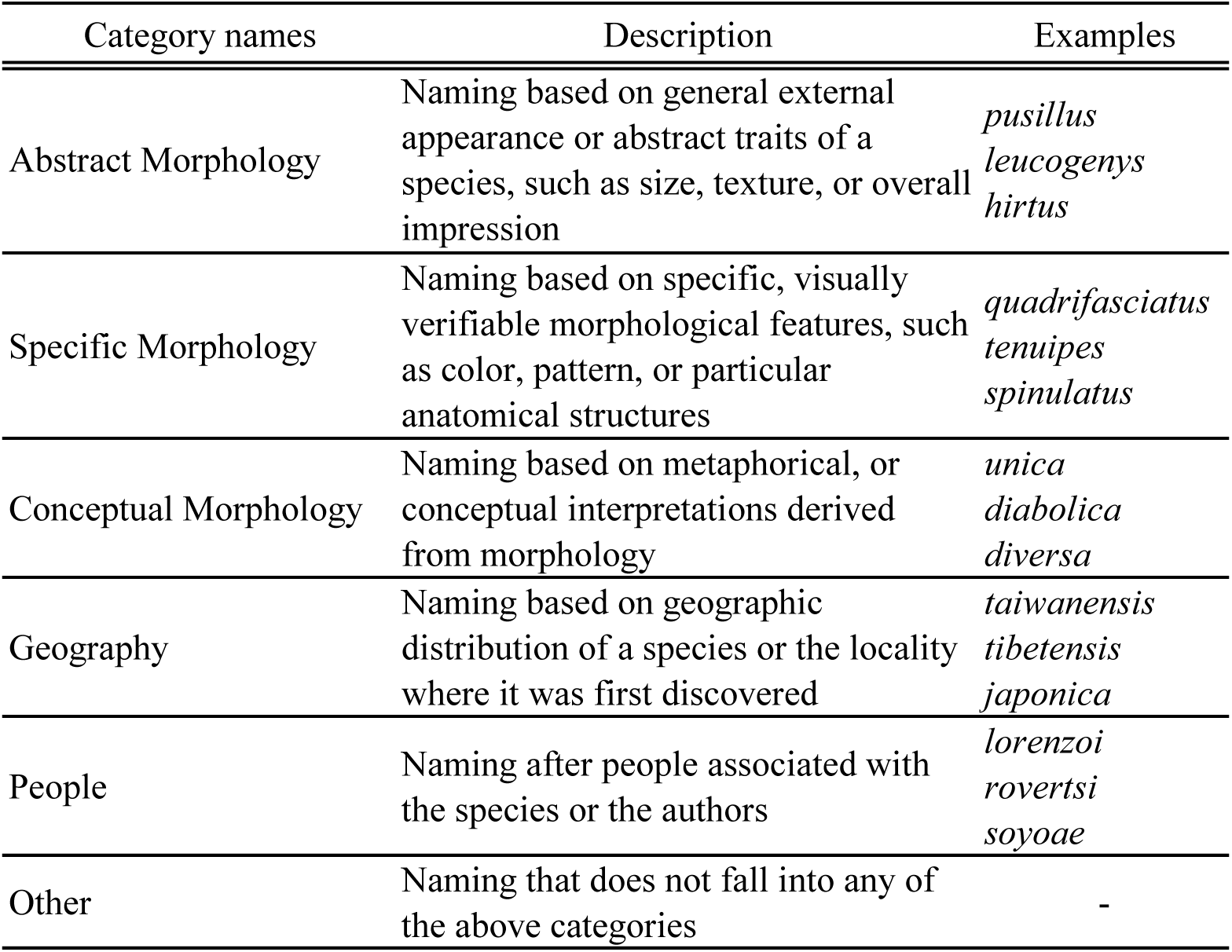
Etymological classification established in this study, defining the six categories used throughout the analyses.

To assess whether these labels generalise beyond spiders, we extracted a random subset of 5,000 species from the Animalia dataset and performed principal component analysis (PCA) on the embeddings. In the PCA scatter plot, epithets assigned to the same category formed distinct and coherent clusters, indicating that the categories derived from spiders are applicable to epithets across Animalia (Figure S2).

### (c) LLM-based inferences

For the full Animalia dataset, we assigned each epithet to one or more etymology categories using an LLM (Table S1). The model was prompted to return all applicable labels that we established: *Abstract Morphology*, *Specific Morphology*, *Conceptual Morphology*, *Geography*, *People* and *Other*. The detailed prompt used for this task has been described in previous work [5], and is therefore omitted.

Initially, the LLM was instructed to distinguish between *Male* and *Female* categories in order to capture potential gender bias in eponymous naming. However, the number of epithets classified as *Female* was extremely small relative to *Male* across the dataset. To avoid instability in downstream analyses and to focus on broader naming patterns, we therefore merged these two labels into a single *People* category in all subsequent analyses.

In addition, we applied an established LLM-based approach to infer the likely nationality of authors [28,29]. For each author string in the Animalia dataset, an LLM was prompted to estimate the most plausible country of origin based solely on the personal name. The model returned a single country label for each author, which we treated as approximations of nationality. Predicted countries were subsequently assigned to broader cultural regions following the classification scheme; African Names, East Asian Names, European Names, Latin American Names, Middle Eastern Names, and South Asian Names (Table S2) [28]

### (d) Computational environment and model specifications

Preliminary clustering and LLM-based classification were conducted on a MacBook Air (M1, 2020) with 16 GB of RAM. The full inference of etymological labels was executed through the OpenAI API, and therefore local GPU resources were not required in this study. We pre-processed the data in Python 3.11.3 using standard libraries including pandas, numpy, scikit-learn, and matplotlib. For the LLM-based procedures, we used the GPT-4o-mini model accessed via the openai package, with the temperature parameter set to 0 and max_tokens fixed at 150 to obtain stable and repeatable outputs.

### (e) Statistical analysis and visualisation

All statistical analyses were conducted using R version 4.4.1 [30] with the packages dplyr, tidyr, mgcv, ggplot2 and pheatmap.

To describe temporal trends in naming, we first calculated annual counts of species for each category. For the Animalia dataset and for each phylum separately, we summed the number of species whose epithet was assigned to a given category in each year.

Several phyla were excluded from the analysis due to small sample sizes (fewer than 250 species), which precluded meaningful visualization of temporal patterns. These phyla include Chaetognatha, Ctenophora, Cycliophora, Dicyemida, Entoprocta, Gnathostomulida, Hemichordata, Loricifera, Micrognathozoa, Orthonectida, Phoronida, Placozoa, Priapulida, and Sipuncula.

Then we modelled temporal changes of naming trends in the proportion of epithets belonging to each category using generalised additive models (GAMs). For each year-category combination, we computed the total number of species (denominator) and the number of species with presence of that category (numerator), yielding an annual proportion. We fitted binomial GAMs with a quasibinomial error distribution and logit link using mgcv::gam, of the form

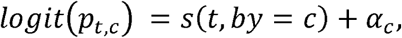

where *p_t,C_* is the proportion of species in year *t* belonging to category *c, s*(*t, by = c*) is a category-specific smooth function of year, and α_*C*_ is a categorical factor representing the etymology category. Annual sample sizes were used as weights. For each model, we obtained fitted proportions on the response scale and associated 95% confidence intervals. In addition, Cycliophora, Micrognathozoa, Orthonectida, Phoronida, Placozoa, and Priapulida were excluded from the GAM analyses owing to insufficient temporal coverage and/or sample size (fewer than 30 species), which precluded reliable model fitting and visualization.

To summarise differences in etymological category usage across taxonomic and cultural groupings, we generated two heatmaps of categories by phylum and by cultural class. In both cases, species were grouped by phylum or cultural class, and the mean of each binary category indicator was calculated within each group, corresponding to the proportion of species whose epithets were assigned to each category. The phylum-category heatmap was restricted to groups represented by at least 100 species to ensure stable estimates. All heatmaps were generated using unscaled values with a continuous colour gradient to emphasise relative differences in category usage among groups.

Finally, to examine temporal shifts in the cultural composition of the taxonomic community, we summarised, for each year, the number of species described by authors belonging to each cultural region and converted these into proportions.

## 3. Results

### (a) Temporal dynamics of species-name etymology

Figure 1 shows that the annual number of new species descriptions increased after the mid-19th century. The most prolific phase occurred following pronounced declines around the 1920s and the 1950s, a pattern visible across almost all phyla. At the Animalia-wide level, all categories increased in absolute frequency.

**Figure 1.**
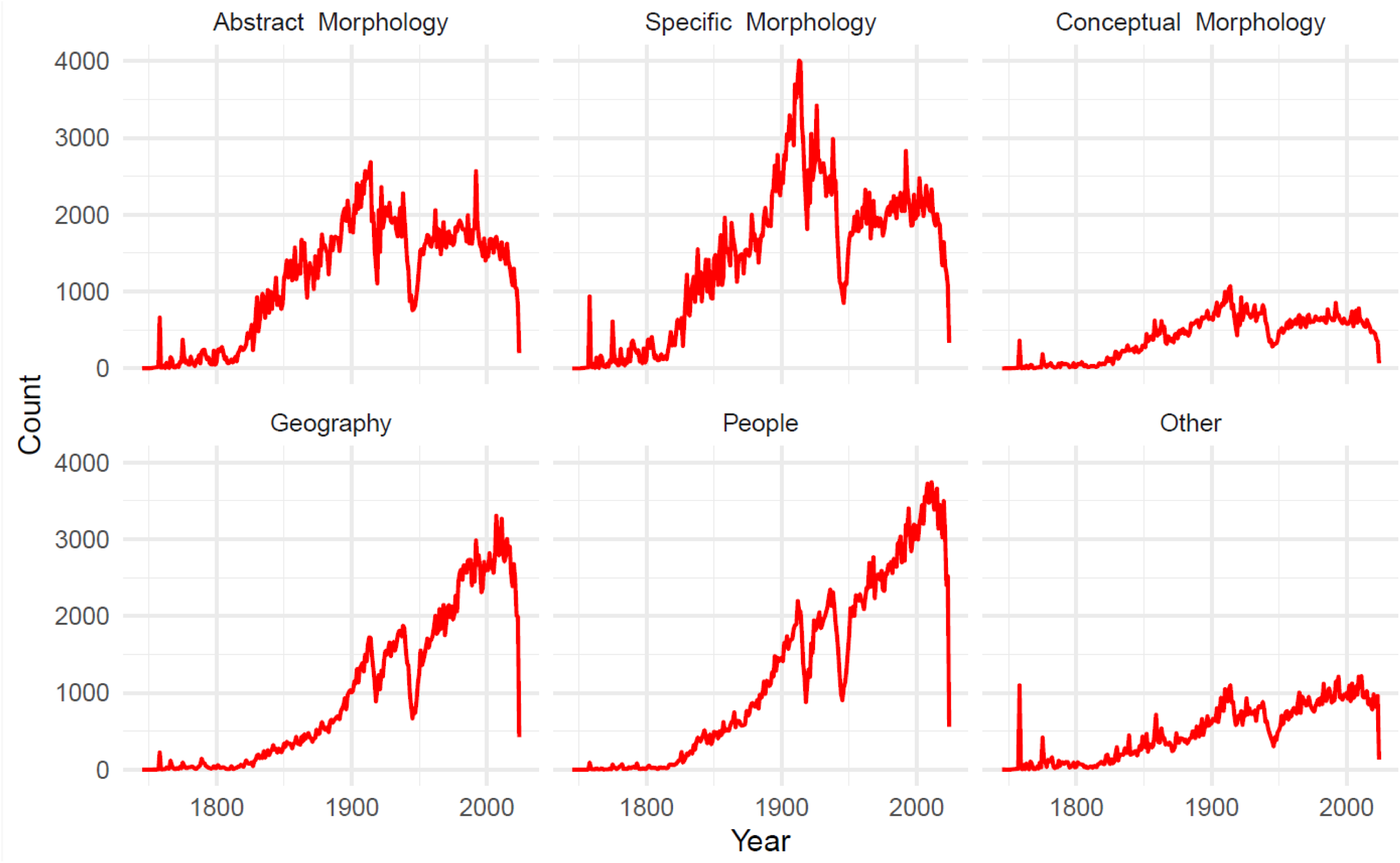
Temporal changes in the annual number of Animalia species assigned to each etymological category, illustrating long-term shifts in naming practices.

Temporal trends in major phyla are shown in Figure 2. Arthropoda closely mirrored the Animalia-wide trends. However, some other phyla exhibited distinct temporal trajectories. Acanthocephala, Bryozoa, Chordata, Cnidaria, Echinodermata, and Rotifera displayed relatively stable temporal trajectories. On the other hand, Kinorhyncha showed a pronounced increase in *People* in the early 21st century.

**Figure 2.**
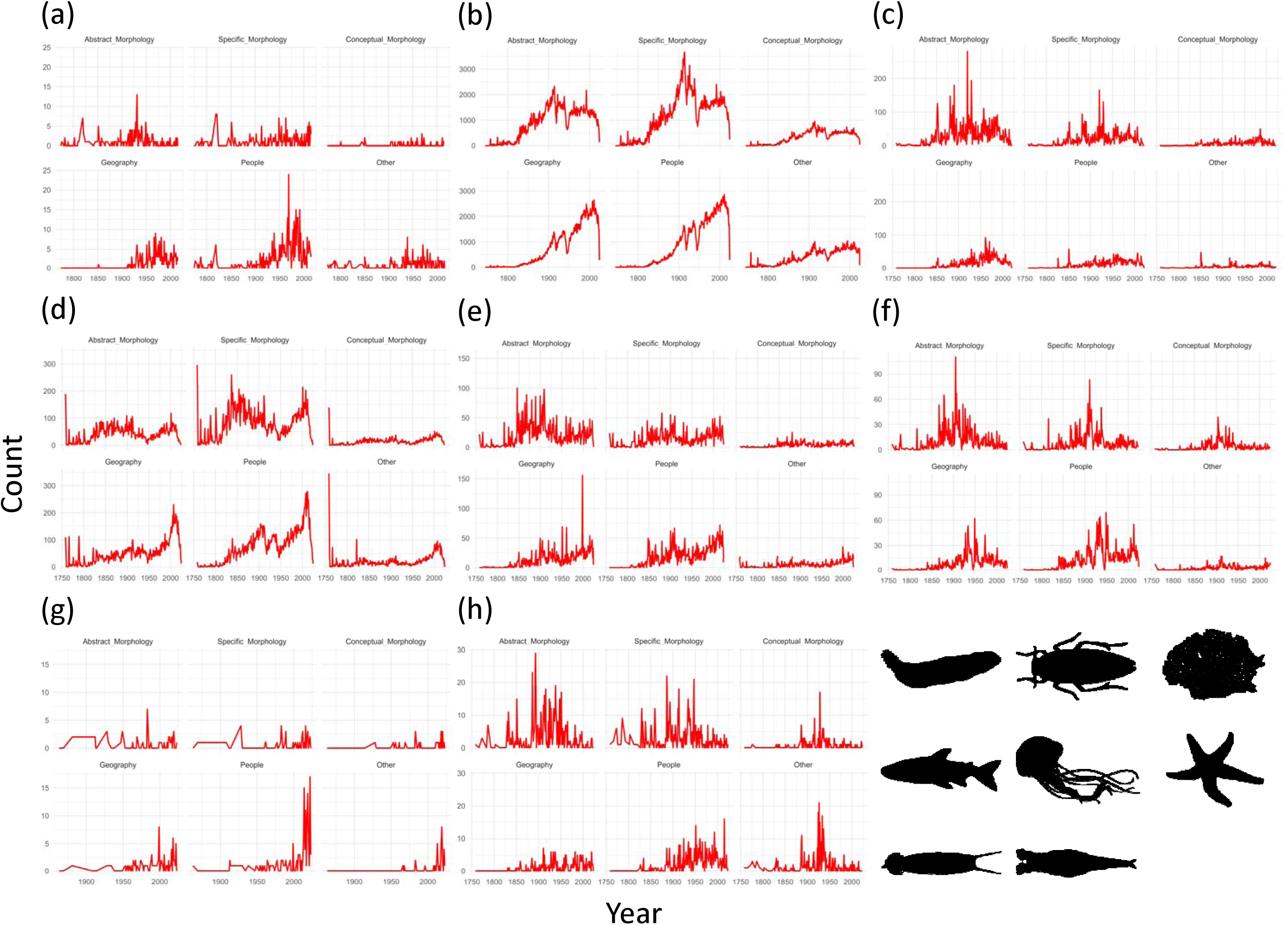
Etymological trends in the total number of species classified into each category for selected phyla: (a) Acanthocephala. (b) Arthropoda. (c) Bryozoa. (d) Chordata. (e) Cnidaria. (f) Echinodermata. (g) Kinorhyncha. (h) Rotifera.

### (b) Shifts in GAM proportion of etymological categories over time

GAMs revealed long-term shifts in the relative frequencies of etymological categories, complementing the absolute temporal dynamics described above, as shown in Figure 3. At the Animalia level, morphology-based epithets (*Abstract Morphology*, *Specific Morphology*, and *Conceptual Morphology*) declined across the past two centuries, while *Geography* and *People* increased continuously, becoming dominant components of modern naming practices.

**Figure 3.**
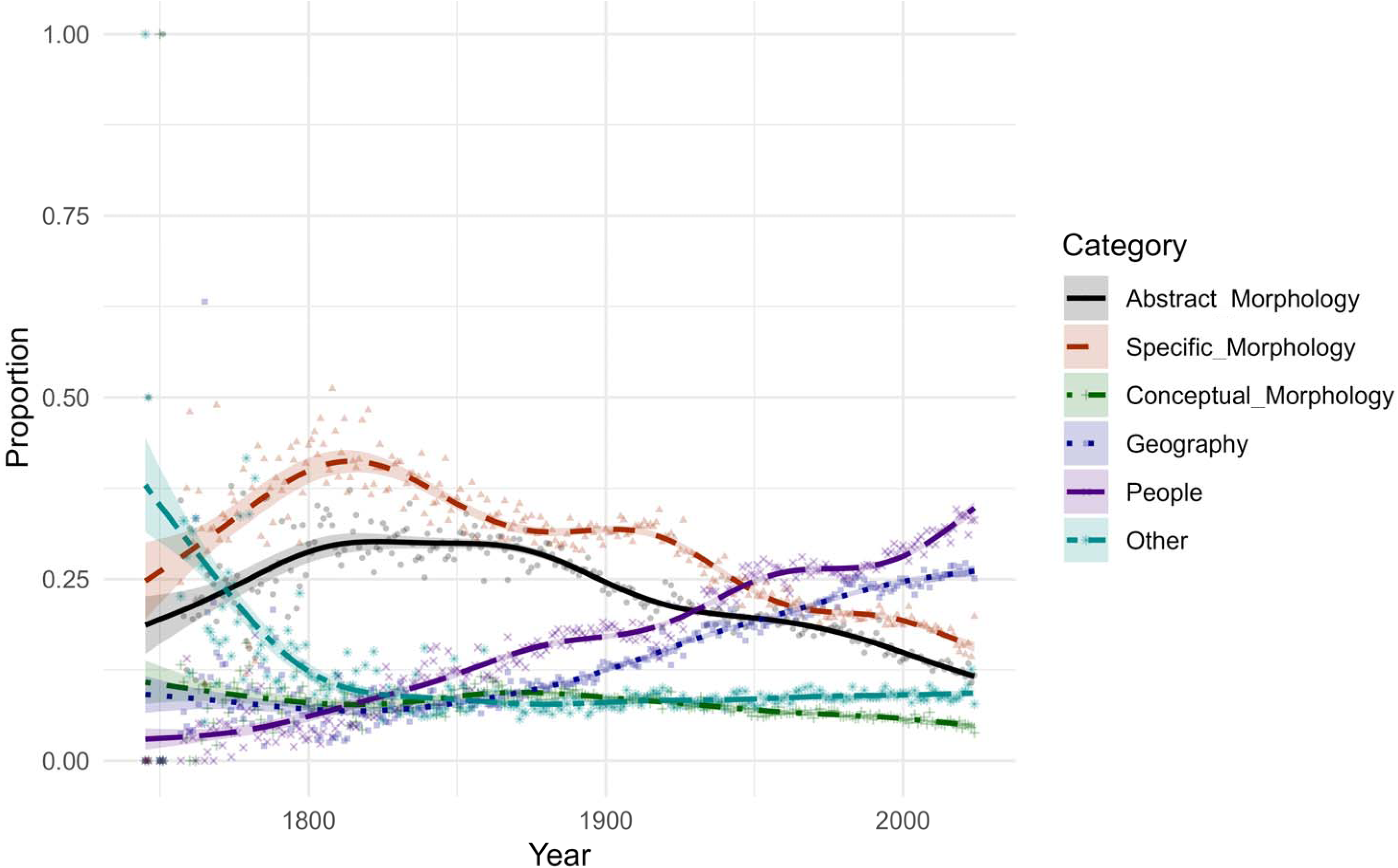
Temporal shifts in the proportional use of the categories estimated using GAMs. Dots represent annual observations, and smooth lines depict GAM-fitted trends with 95% confidence intervals. Categories are coded by colour and line-type as follows: *Abstract Morphology* (solid, black), *Specific Morphology* (dashed, dark red), *Conceptual Morphology* (dot-dashed, dark green), *Geography* (dotted, dark blue), *People* (long-dashed, purple), and *Other* (two-dashed, light blue).

Several phyla exhibited distinct temporal trajectories, whereas Arthropoda showed GAM patterns nearly identical to those of Animalia as a whole. (Figure 4). In contrast to the general trends, Nematoda exhibited a declining use of *People* beginning around the early 2000s, highlighting a recent shift in naming conventions in this group. The *People* epithets in Annelida and Porifera increased around 2000. Bryozoa illustrated the increase not only in *People* but also in *Geography* after 2000. In Chordata, the GAMs showed a rise in *People* throughout the 19th century and a sharp increase of *Geography* around 2000.

**Figure 4.**
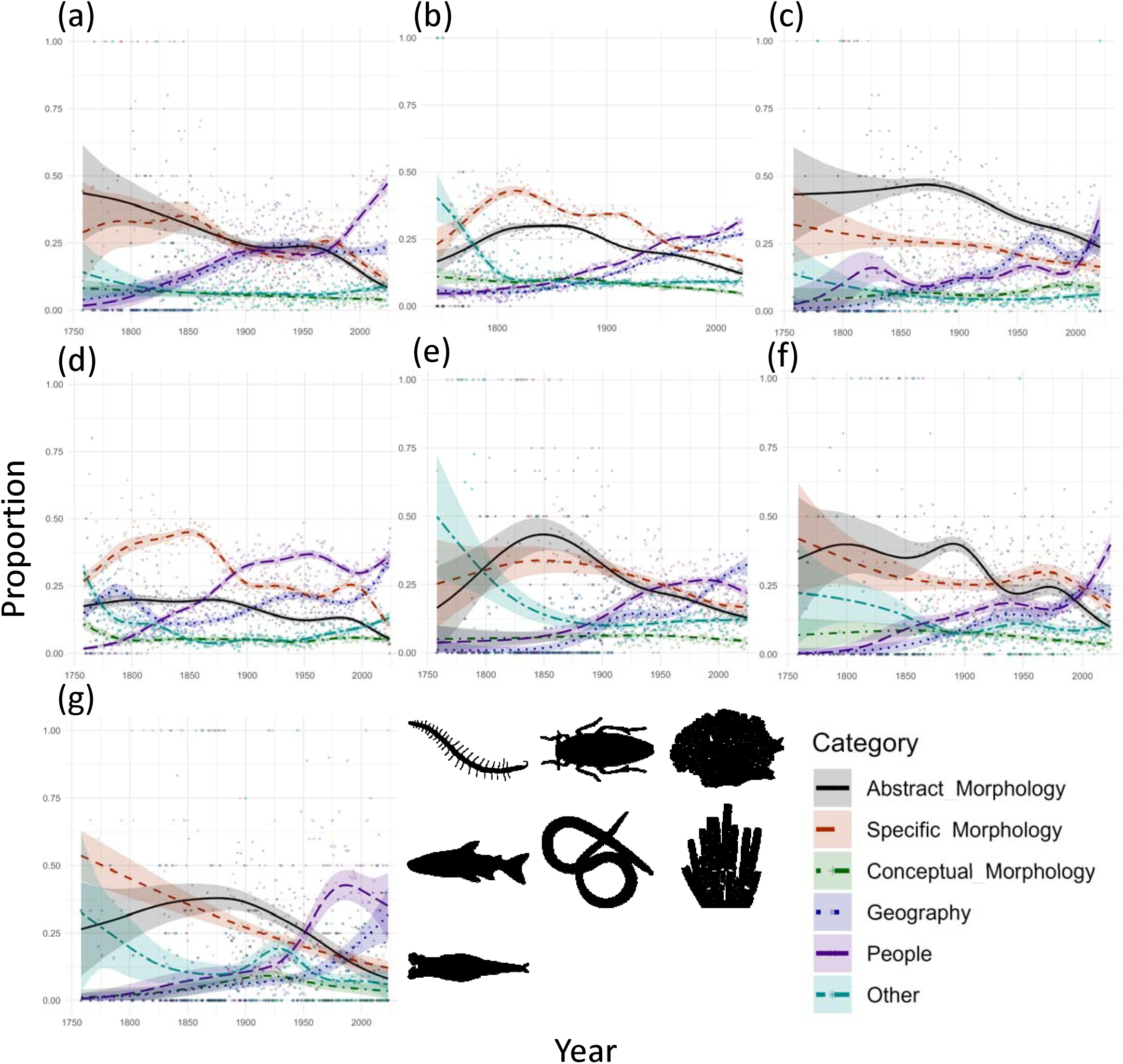
Temporal shifts in the proportional use of etymological categories for selected phyla, shown using the same colour and line-type scheme and GAM settings as in Figure 3: (a) Annelida. (b) Arthropoda. (c) Bryozoa. (d) Chordata. (e) Nematoda. (f) Porifera. (g) Rotifera.

Rotifera displayed yet another unique trend. While *People* increased beginning around the 1950s, *Specific Morphology* showed a persistent and monotonic decline across the entire timespan. These temporal shifts are distinct in both timing and magnitude compared with other phyla.

### (c) Inter-phylum differences in naming practices

Marked differences in naming conventions emerged across phyla, revealing strong lineage-specific traditions in the use of etymological categories (Figure 5).

**Figure 5.**
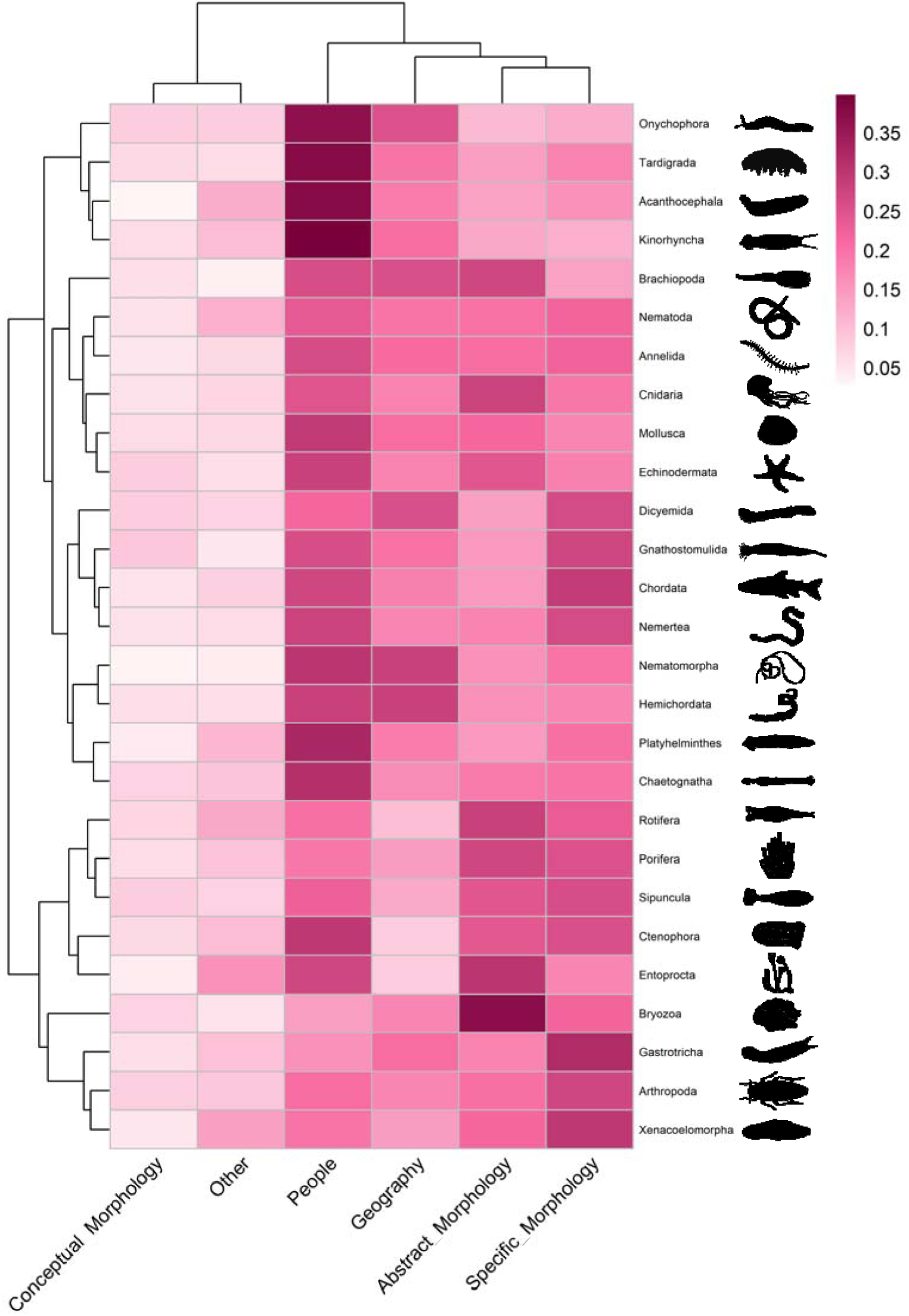
Heatmap showing the relative frequency of etymological categories across Animalia. Rows represent phyla and columns represent the categories. Cell colours indicate the relative frequency of each category within a phylum. Rows and columns are ordered by hierarchical clustering based on similarity in category composition. Representative silhouettes of each phylum are shown on the right.

Acanthocephala, Kinorhyncha, Onychophora, and Tardigrada showed some of the highest proportional use of *People* among all phyla.

Although *People* was the most common naming category across Animalia, several phyla showed deviations from the global pattern. In Arthropoda, Bryozoa, Dicyemida, Gastrotricha, Porifera, Rotifera, Sipuncula and Xenacoelomorpha, People-based epithets were used less relative to other groups. While *Abstract Morphology* is frequently used in Bryozoa to compensate for low prevalence of *People*, Arthropoda, Bryozoa, Gastrotricha, and Xenacoelomorpha exhibited a notably high proportion of *Specific Morphology*. In addition, the proportional use of *Geography* was low in Ctenophora, Entoprocta, Porifera, Rotifera, and Sipuncula.

### (d) Geographic and cultural influences on taxonomy

Naming conventions varied among regions defined by inferred nationalities of authors (Figure 6). Latin American authors exhibited the highest proportional use of *People*, followed by Middle Eastern and South Asian authors. In all three regions, *Geography* represented the second-most common category. In contrast, East Asian authors showed remarkably lower proportional use of *People*. The epithets described by East Asian authors were characterised by a predominance of *Geography* over *Specific Morphology*.

**Figure 6.**
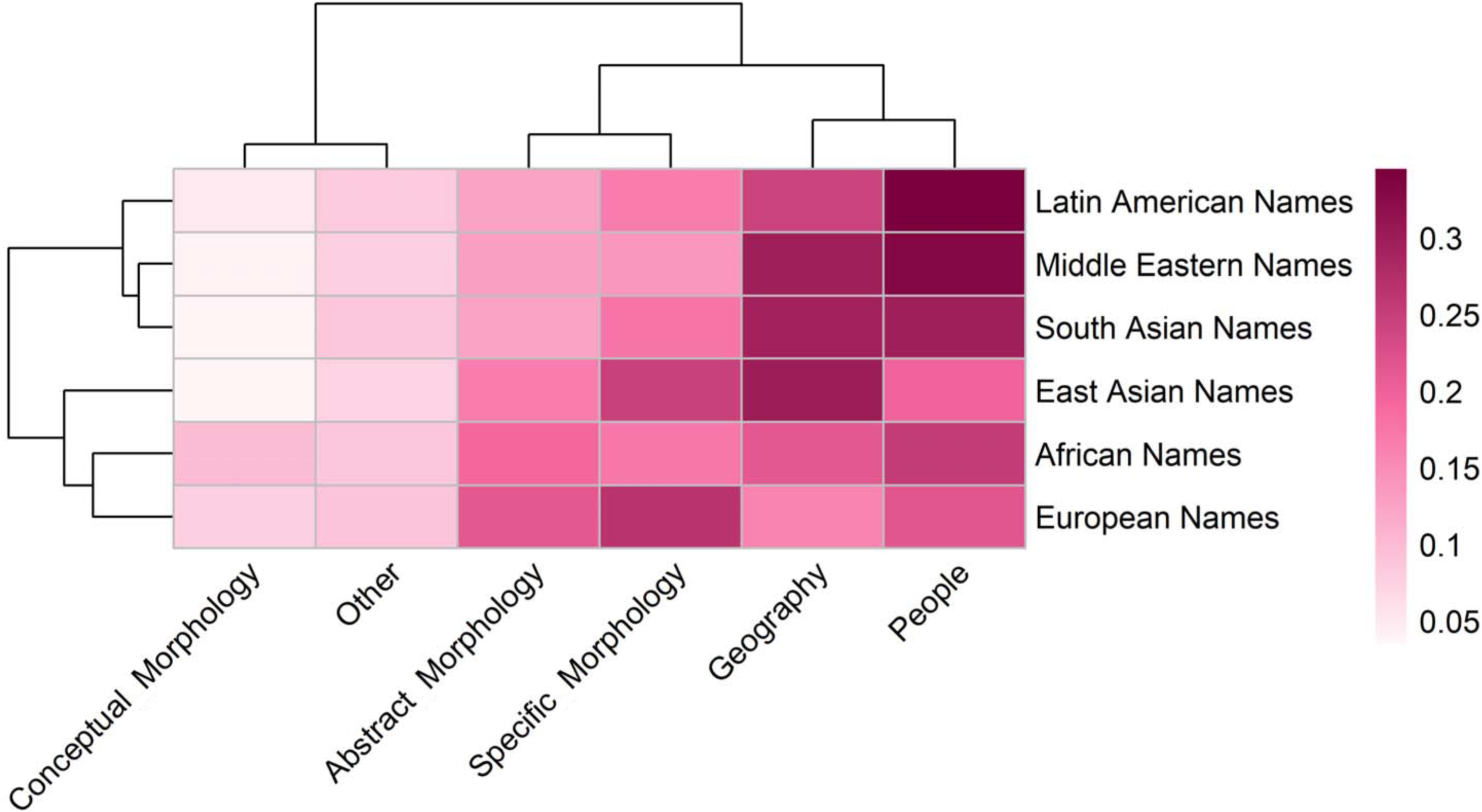
Heatmap illustrating the association between etymological categories and cultural name classes. Rows represent cultural name classes and columns represent the categories. Cell colours indicate the relative frequency of each category within a cultural class. Dendrograms show hierarchical clustering based on similarity in etymological composition.

European authors showed a distinct pattern. Although their use of *People* is relatively low, second only to East Asia, they also show sparse use of *Geography*, which was uniquely low compared with all other regions analysed. Instead, *Specific Morphology* was the most common category.

Beyond differences in naming categories, the cultural composition of the taxonomist community itself has shifted markedly over time (Figure 7). Until the early 20th century, nearly all species were described by scientists of inferred European origin. However, beginning in the mid-20th century, authors from other regions entered the taxonomy. Their relative contributions grew most prominently after the 1970s, producing a gradual but detectable diversification in the authorship.

**Figure 7.**
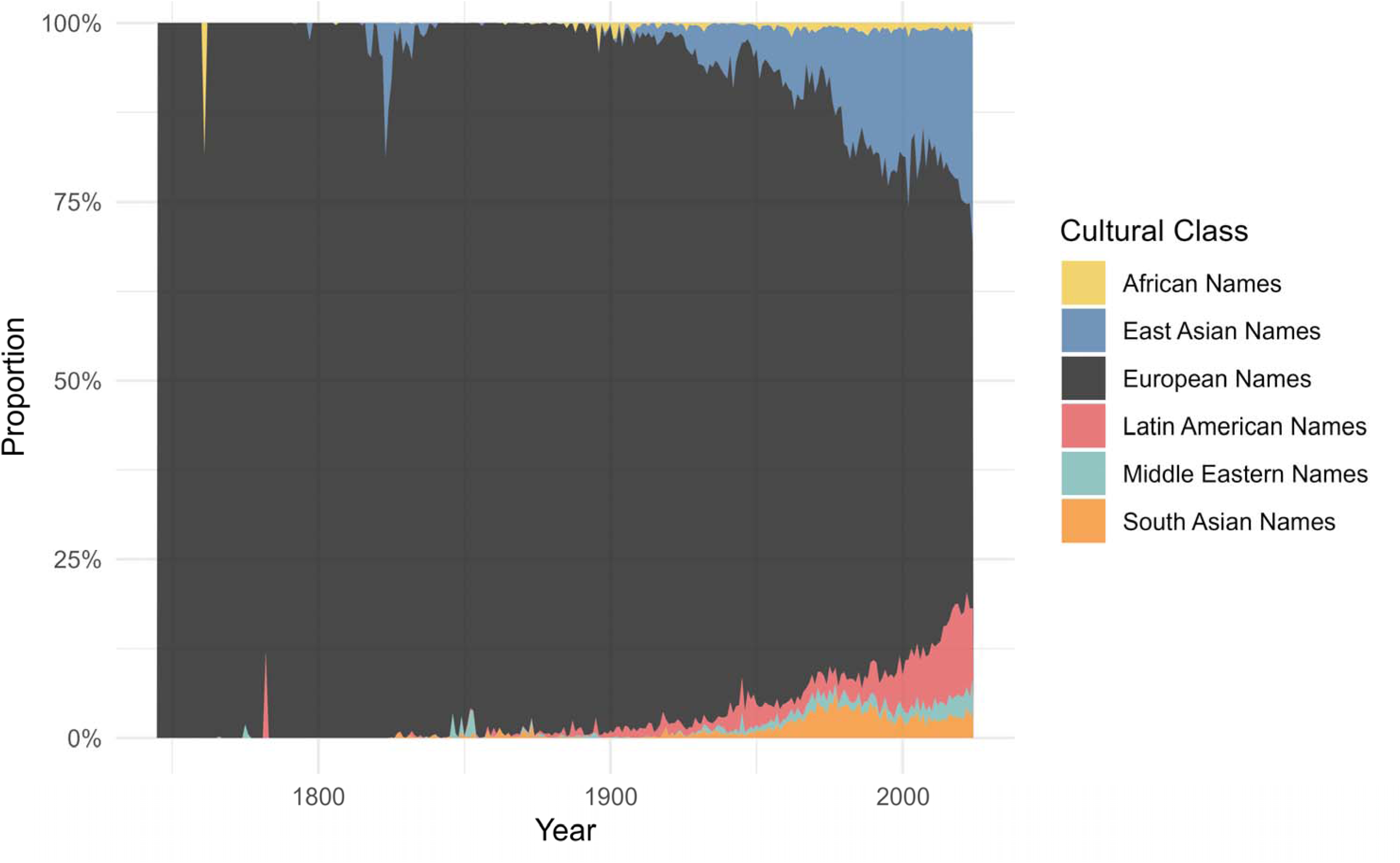
Temporal changes in the proportional composition of cultural name classes across Animalia. Stacked areas represent the relative contribution of each cultural class to all species named in a given year, with proportions summing to 100%.

## 4. Discussion

Our results reveal clear historical, taxonomic, and cultural structure in the etymology of animals’ scientific names. Taken together, these patterns show that naming practices, while constrained by the formal rules of the Code [1], have been shaped by scientific traditions, methodological developments, and the global expansion of the taxonomist community. Below, we discuss major aspects of these findings.

### (a) Historical factors

Our results reveal that large-scale historical events have left clear signatures on the trajectory of species naming. The pronounced declines in description around the 1920s and the 1950s (Figure 1) are likely related to the global disruption caused by World War I (1914-1918) and World War II (1939-1945). During this period, taxonomic activity was widely deprioritised as scientific personnel and resources were directed toward wartime efforts [31]. The sharpness of this decline varies across phyla (Figure 2): groups with long-standing, stable research attention such as Chordata, Cnidaria, Echinodermata, and Rotifera exhibited comparably muted declines. This pattern does not necessarily indicate post-war recovery of description rates; rather, they may have been well studied consistently throughout their taxonomic history as relatively familiar groups.

### (b) Taxonomic and methodological factors

Temporal patterns observed across Animalia were strongly dominated by Arthropoda, which comprise 1,193,760 species of the 1,538,390 species in our dataset, accounting for nearly 80% of all described animal species analysed here. As a consequence, apparent Animalia-level trends primarily reflect the dynamics of Arthropoda rather than patterns that are uniformly shared across phyla. This abundance of Arthropoda is consistent with previous estimates that they constitute the vast majority of animal biodiversity [32]. This underscores the importance of phylum-level comparisons, as global patterns alone mask substantial lineage-specific trajectories.

Across Animalia, the decline of morphology-based epithets (*Abstract Morphology*, *Specific Morphology*, and *Conceptual Morphology*) and the rise of *Geography* and *People* are the most prominent trends (Figure 3) consistent with previous analyses [3,8,26,33]. One plausible explanation is that the improvements in optical microscopy, the invention of electron microscopes, and molecular techniques (e.g., Polymerase Chain Reaction: PCR, DNA sequencing) enabled taxonomists to subdivide morphologically similar species. As species descriptions have increasingly relied on subtle diagnostic differences not easily expressible as epithets, names referencing localities or people may have become more practical and preferred. In this context, the historical increase of *People* epithets cannot be viewed independently of contemporary debates surrounding eponyms, including calls for renaming and proposals for revision of the Code [12,14–18]. Understanding the long-term drivers behind the prevalence of eponyms in taxonomy is therefore essential for interpreting current discussions on nomenclatural stability and reform.

Taxon-specific histories also shape naming traditions (Figure 4). The persistent monotonic decline of *Specific Morphology* in Rotifera may reflect the difficulty of expressing morphological differences in extremely small, morphologically conservative organisms. Bryozoa exhibits a sharp post-2000 increase in *People* and *Geography*, likely reflecting the influence of a narrow taxonomist community. In this phylum, although 1,680 species descriptions are recorded after 2000, only 260 distinct author patterns are represented. This indicates that species descriptions are concentrated among a relatively small number of taxonomists, and consequently, the naming preferences of these few taxonomists are disproportionately reflected in the overall etymological patterns observed in this taxon.

In several marine-dominated phyla, the elevated frequency of *People* may reflect long-standing naming conventions tied to oceanographic research. Marine taxonomy has a tradition of commemorating research vessels involved in specimen collection (e.g., *Galacantha valdiviae Balss,* 1913; *Parapagurus shibogae* de Saint Laurent, 1972) [34,35]. Because the names of ships are often treated as feminine nouns, this may contribute to the increased use of *People*. This pattern indicates that naming practices may arise not only from the preferences of individual taxonomists but also from the history of biological fieldwork itself.

Importantly, we did not use an etymological label that refers to host-related epithets used for parasitic animals. Therefore, the recent decline of *People* in Nematoda (Figure 4) may reflect a pattern specific to parasitic groups that some epithets might be misclassified into our categories. However, our goal in this study was to identify a broad Animalia-wide naming pattern. These deviations are unlikely to affect the interpretation of the results.

### (c) Cultural and geographic factors

The inferred origins of authors reveal a pronounced cultural structure of naming practices (Figure 5, 6). European authors showed high proportional use of *Specific Morphology* and notably low use of *Geography* and *People*. This pattern is consistent with the historical roots of taxonomy in European scientific traditions. Early naturalists, working during the colonial expansion, described large numbers of familiar organisms, species of societal importance, or morphologically distinctive taxa in the 18th and 19th centuries. This background led to an etymological record strongly shaped by descriptive and honorific practices rooted in Europe.

In contrast, authors from Latin America, the Middle East, and South Asia exhibited much higher use of *People* and *Geography*. These patterns may reflect a greater emphasis on recognising their colleagues or regions. It is likely that the taxonomist communities outside Europe are developing their own naming cultures as global participation has expanded.

East Asian authors showed particularly low use of *People*. One hypothesis for this pattern is that relatively short and structurally similar surnames common in some East Asian countries such as China and Korea may limit their suitability [36–38]. It might lead to low frequencies of naming for colleagues or authority in their own communities. The pattern highlights that cultural and linguistic structures can also affect scientific naming practices.

A low proportional use of *Geography* was observed in Ctenophora, Entoprocta, Porifera, Rotifera, and Sipuncula. This pattern may be related to the high proportion of species named by European researchers; however, this cannot be concluded from the present study alone.

Although authorship has diversified substantially since the mid-20th century with Latin America and East Asia showing pronounced growth, European authors still contribute the majority of species descriptions (Figure 7). The underrepresentation of Africa, the Middle East, and South Asia likely reflects global asymmetries in scientific infrastructure and funding [39], which continue to shape both who names species and how species are named.

Even though the LLM-based inference of authors’ nationality has high accuracy, these assignments should be interpreted as coarse cultural proxies rather than precise national identities.

### (d) Implications for understanding the transition in zoological nomenclature

Our study demonstrates that species names encode not only biological information but also the historical, cultural, and methodological contexts. Because naming practices arise from the intersection of scientific norms, community structure, linguistic constraints, geography, and political history, understanding the transition requires considering all these dimensions together.

The pronounced variation among phyla and regions shows that zoological nomenclature is not a uniform system but a culturally embedded practice shaped by who participates in taxonomy and how they engage with organisms.

## Authors’ contributions

KN designed the study, performed the data analyses, and wrote the manuscript. KI developed the LLM prompt and prepared the data for analyses. KI, HS, and TT contributed substantially to improving the manuscript drafts. All authors approved the final version of the manuscript.

## Acknowledgements.

We acknowledge the Catalogue of Life (https://www.catalogueoflife.org/) for providing comprehensive, open-access taxonomic data available. Silhouette images were obtained from PhyloPic (https://www.phylopic.org/). We thank Jun Togashi for sharing valuable comments on the manuscript, and Takashi Yoshida for kindly facilitating the exchange and discussion.

## Supplementary materials

Table S1. Summary of etymological category composition across Animalia and individual phyla. The table reports the total number of species names, the number of species assigned to each category, and the corresponding proportions. Because category assignments are non-exclusive, a single species name may contribute to multiple categories; therefore, category totals can exceed the total number of species.

Table S2. Country-level mapping of cultural name classes used in this study. Each row represents a single country and its assigned cultural name class, based on dominant linguistic and historical traditions. Countries with predominantly European settler origins (e.g., the United States, Canada, Australia, and New Zealand) were grouped within European Names, reflecting the linguistic and cultural origins of scientific naming practice rather than present-day geography. North African Countries (e.g., Algeria, Egypt, Libya, Morocco, and Tunisia) were assigned to Middle Eastern Names based on shared Arabis linguistic and cultural traditions. Fiji and Papua New Guinea were treated separately from continental Asian categories owing to their distinct Melanesian linguistic traditions.

Figure S1. Average silhouette scores for K-means clustering in spider’s dataset across different values of k (6-11). The highest score was observed at k =7.

Figure S2. Two-dimensional t-SNE of ICA-reduced semantic embedding in the spider’s dataset, coloured by K-means cluster assignment (k = 7).

Figure S3. Semantic structure of Animalia epithets in PCA plot. Each point represents a single species epithet, positioned according to its semantic similarity to others. Points are coloured by etymological category, including single-category assignments and combinations of multiple categories. To aid visual interpretation of the three-dimensional structure, the same point cloud is shown from multiple viewing directions. The schematic cube (lower right) indicates the viewing directions.

Figure S4. Etymological trends in the total count of species classified into each category for additional phyla not used for interpretation in the main text: (a) Annelida. (b) Brachiopoda. (c) Gastrotricha. (d) Mollusca. (e) Nematoda. (f) Nematomorpha. (g) Nemertea. (h) Onychophora. (i) Platyhelminthes. (j) Porifera. (k) Tardigrada. (l) Xenacoelomorpha.

Figure S5. Temporal trends in the proportional use of etymological categories for phyla not shown in Figure 4., using the same colour and line-type scheme and GAM settings as in Figure 3: (a) Acanthocephala. (b) Brachiopoda. (c) Chaetognatha. (d) Cnidaria. (e) Ctenophora. (f) Dicyemida. (g) Echinodermata. (h) Entoprocta. (i) Gastrotricha. (j) Gnathostomulida. (k) Hemichordata. (l) Kinorhyncha. (m) Loricifera. (n) Mollusca. (o) Nematomorpha. (p) Nemertea. (q) Onychophora. (r) Platyhelminthes. (s) Sipuncula. (t) Tardigrada. (u) Xenacoelomorpha.

## References

1. International Commission on Zoological Nomenclature. 1999 International Code of Zoological Nomenclature. 4th ed. London: International Trust for Zoological Nomenclature. (doi:10.5962/bhl.title.50608)

2. Jasper PD, Froehlich EM, Carbayo-Baz FJ. 2021 A study on the etymology of the scientific names given to planarians (Platyhelminthes, Tricladida) by Ernest Marcus’ school. Papéis Avulsos de Zoologia 61, e20216105. (doi:10.11606/1807-0205/2021.61.05)

3. Nojiri K, Inoshita K, Sugeno H. 2025 Automated labeling of scientific names and etymological trend analysis in phytophagous arthropods using large language model. jzoo 42, 492–497. (doi:10.2108/zs250025)

4. Mlynarek JJ, Cull C, Parachnowitsch AL, Vickruck JL, Heard SB. 2023 Can species naming drive scientific attention? A perspective from plant-feeding arthropods. Proceedings of the Royal Society B: Biological Sciences 290, 20222187. (doi:10.1098/rspb.2022.2187)

5. Inoshita K, Nojiri K, Sugeno H, Taga T. 2025 Evaluation of the automated labeling method for taxonomic nomenclature through prompt-optimized large language model. 2025 IEEE International Conference on Industry 4.0, Artificial Intelligence, and Communications Technology (IAICT), 528–535. (doi:10.1109/IAICT65714.2025.11100523)

6. Macêdo RL, Elmoor-Loureiro LMA, Sousa FDR, Rietzler AC, Perbiche-Neves G, Rocha O. 2023 From pioneers to modern-day taxonomists: the good, the bad, and the idiosyncrasies in choosing species epithets of rotifers and microcrustaceans. Hydrobiologia 850, 4271–4282. (doi:10.1007/s10750-023-05302-7)

7. Vendetti J. 2022 Gender representation in molluscan eponyms: Disparities and legacy. malb 39, 19–31. (doi:10.4003/006.039.0106)

8. Mammola S, Viel N, Amiar D, Mani A, Hervé C, Heard SB, Fontaneto D, Pétillon J. 2023 Taxonomic practice, creativity and fashion: What’s in a spider name? Zoological Journal of the Linnean Society 198, 494–508. (doi:10.1093/zoolinnean/zlac097)

9. Figueiredo E, Smith GF. 2010 What’s in a name: epithets in *Aloe* L. (Asphodelaceae) and what to call the next new species. brad 2010, 79–102. (doi:10.25223/brad.n28.2010.a9)

10. Poulin R, McDougall C, Presswell B. 2022 What’s in a name? Taxonomic and gender biases in the etymology of new species names. Proceedings of the Royal Society B: Biological Sciences 289, 20212708. (doi:10.1098/rspb.2021.2708)

11. Jozwiak P, Rewicz T, Pabis K. 2015 Taxonomic etymology – in search of inspiration. ZooKeys 513, 143–160. (doi:10.3897/zookeys.513.9873)

12. Pillon Y. 2021 The inequity of species names: The flora of New Caledonia as a case study. Biological Conservation 253, 108934. (doi:10.1016/j.biocon.2020.108934)

13. Schiebinger LL. 1993 Nature’s Body: Gender in the Making of Modern Science. Boston: Beacon Press.

14. Guedes P et al. 2023 Eponyms have no place in 21st-century biological nomenclature. Nat Ecol Evol 7, 1157–1160. (doi:10.1038/s41559-023-02022-y)

15. Rummy P, Rummy JT. 2021 Recontextualising the style of naming in nomenclature. Humanit Soc Sci Commun 8, 283. (doi:10.1057/s41599-021-00975-8)

16. Wright SD, Gillman LN. 2022 Replacing current nomenclature with pre-existing indigenous names in algae, fungi and plants. TAXON 71, 6–10. (doi:10.1002/tax.12599)

17. Gillman LN, Wright SD. 2020 Restoring indigenous names in taxonomy. Commun Biol 3, 609. (doi:10.1038/s42003-020-01344-y)

18. Roksandic M, Musiba C, Radović P, Lindal J, Wu X-J, Figueiredo E, Smith GF, Roksandic I, Bae CJ. 2023 Change in biological nomenclature is overdue and possible. Nat Ecol Evol 7, 1166–1167. (doi:10.1038/s41559-023-02104-x)

19. Ceríaco LMP et al. 2023 Renaming taxa on ethical grounds threatens nomenclatural stability and scientific communication: Communication from the International Commission on Zoological Nomenclature. Zoological Journal of the Linnean Society 197, 283–286. (doi:10.1093/zoolinnean/zlac107)

20. Orr MC et al. 2023 Inclusive and productive ways forward needed for species-naming conventions. Nat Ecol Evol 7, 1168–1169. (doi:10.1038/s41559-023-02103-y)

21. Jablonski D, Dufresnes C. 2024 Nomenclatural censorship puts biodiversity conservation and taxonomic science at risk. Alytes 41, 1–4.

22. Antonelli A et al. 2023 People-inspired names remain valuable. Nat Ecol Evol 7, 1161–1162. (doi:10.1038/s41559-023-02108-7)

23. Pethiyagoda R. 2023 Policing the scientific lexicon: The new colonialism? Megataxa 10, 20–25. (doi:10.11646/megataxa.10.1.4)

24. Jiménez-Mejías P et al. 2024 Protecting stable biological nomenclatural systems enables universal communication: A collective international appeal. BioScience 74, 467–472. (doi:10.1093/biosci/biae043)

25. Heard SB, Mlynarek JJ. 2023 Naming the menagerie: Creativity, culture and consequences in the formation of scientific names. Proceedings of the Royal Society B: Biological Sciences 290, 20231970. (doi:10.1098/rspb.2023.1970)

26. Sugeno H, Inoshita K, Nojiri K, Taga T. 2025 Introducing large language models to human-based etymological classification in zooplankton., 2025.05.08.652882. (doi:10.1101/2025.05.08.652882)

27. Bánki O et al. 2025 Catalogue of Life (Version 2025-04-10). See 10.48580/dgplc (accessed on 18 April 2025).

28. Phonchai T, Siripong S, Patterson N, Campbell O. 2025 Large language models for zero-shot multicultural name recognition. (doi:10.48550/arXiv.2507.04149)

29. Inoshita K. 2026 Nationality and Region Prediction from Names: A Comparative Study of Neural Models and Large Language Models. (doi:10.48550/arXiv.2601.08692)

30. R Core Team. 2024 R: A Language and environment for statistical computing. R Foundation for Statistical Computing. See https://www.r-project.org/ (accessed on 12 March 2025).

31. Gross DP, Sampat BN. 2023 The World War II crisis innovation model: What was it, and where does it apply? Research Policy 52, 104845. (doi:10.1016/j.respol.2023.104845)

32. May RM. 1988 How Many Species Are There on Earth? Science 241, 1441.

33. Inoshita K, Nojiri K, Sugeno H, Taga T. 2025 Evaluation of the automated labeling method for taxonomic nomenclature through prompt-optimized large language model. *arXiv*, arXiv:2503.10662. (doi:10.48550/arXiv.2503.10662)

34. de Saint Laurent M. 1972 Sur la famille des Parapaguridae Smith, 1882. Description de *Typhlopagurus foresti* gen. nov., sp. nov., et de quinze espèces ou sous-espèces nouvelles de *Parapagurus* Smith (Crustacea, Decapoda). Bijdragen tot deDierkunde 42, 97–123. (doi:10.1163/26660644-04202001)

35. Balss H. 1913 Neue Galatheiden aus der Ausbeute der deutschen Tiefsee-Expedition Valdivia. Zoologischer Anzeiger 41, 221–226.

36. Ruofu D. 1986 Surnames in china. Journal of Chinese Linguistics 14, 315–328.

37. Kim BJ, Park SM. 2005 Distribution of Korean family names. Physica A: Statistical Mechanics and its Applications 347, 683–694. (doi:10.1016/j.physa.2004.08.028)

38. Baek SK, Kiet HAT, Kim BJ. 2007 Family name distributions: Master equation approach. *Phys*. Rev. E 76, 046113. (doi:10.1103/PhysRevE.76.046113)

39. DuBay S, Droguett DHP, Piland NC. 2022 Global inequity in scientific names and who they honor., 2020.08.09.243238. (doi:10.1101/2020.08.09.243238)

