## supplementary figures for "Unravelling historical, taxonomic, and cultural influences on the etymology of scientific names across Animalia"

### Slide 1
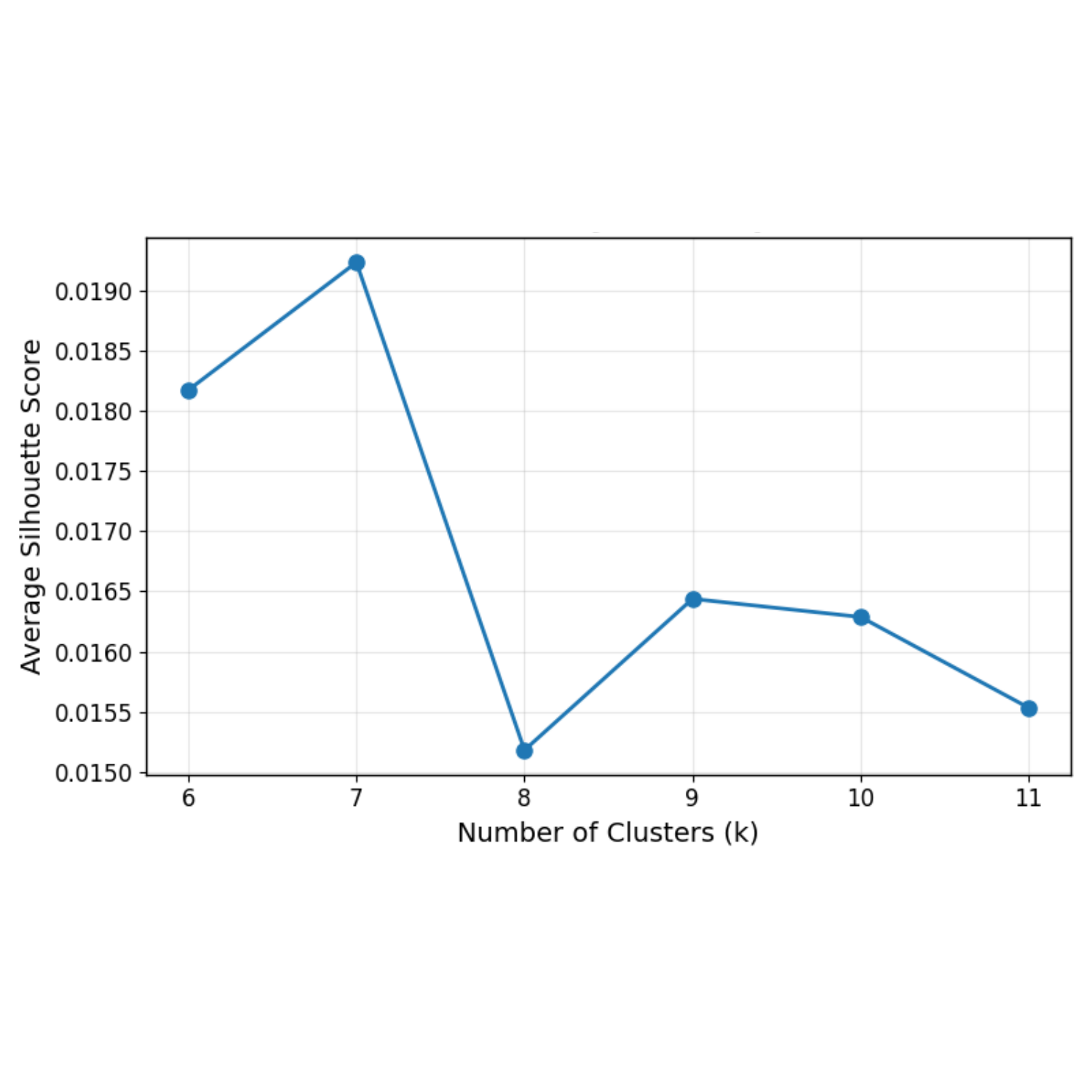

### Slide 2
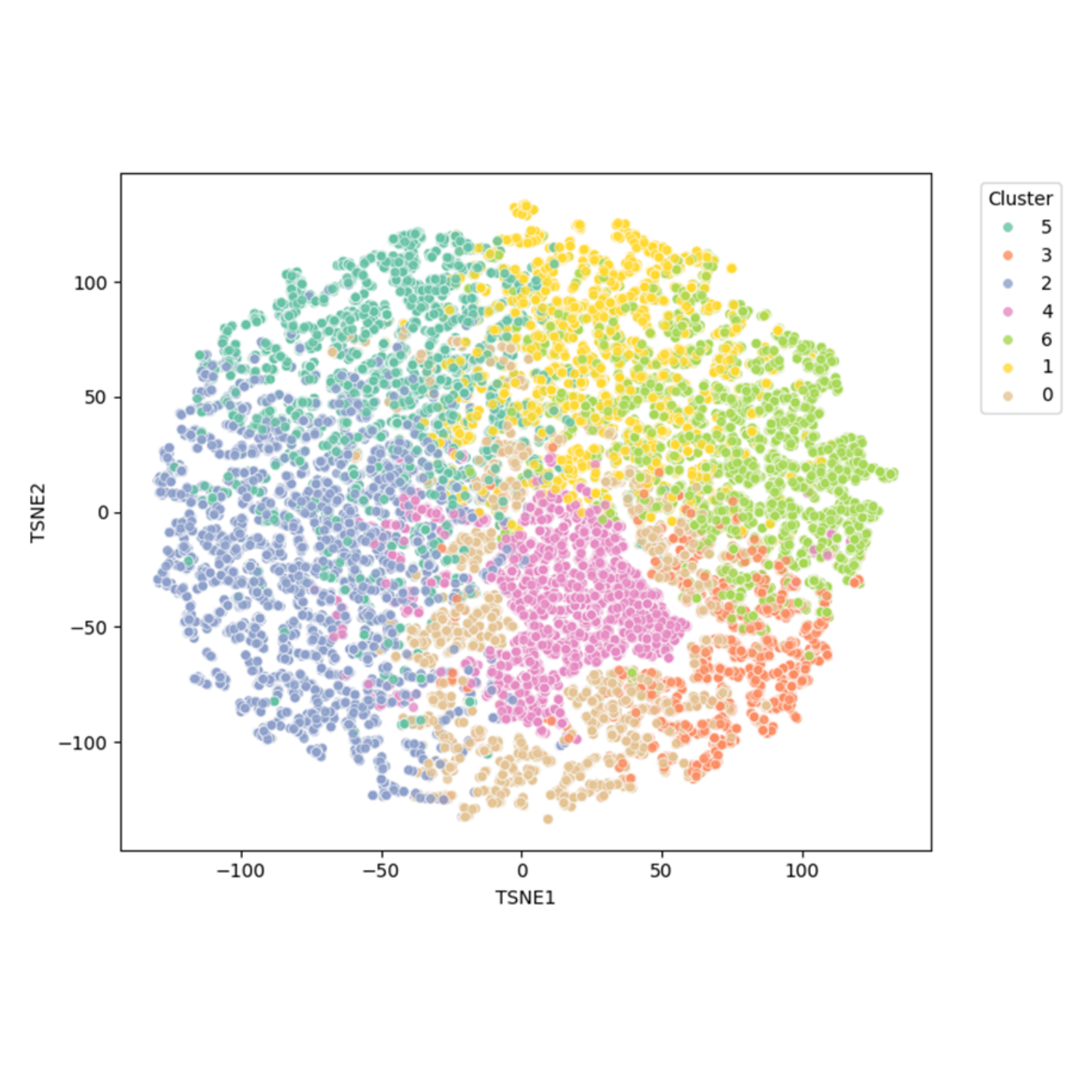

### Slide 3
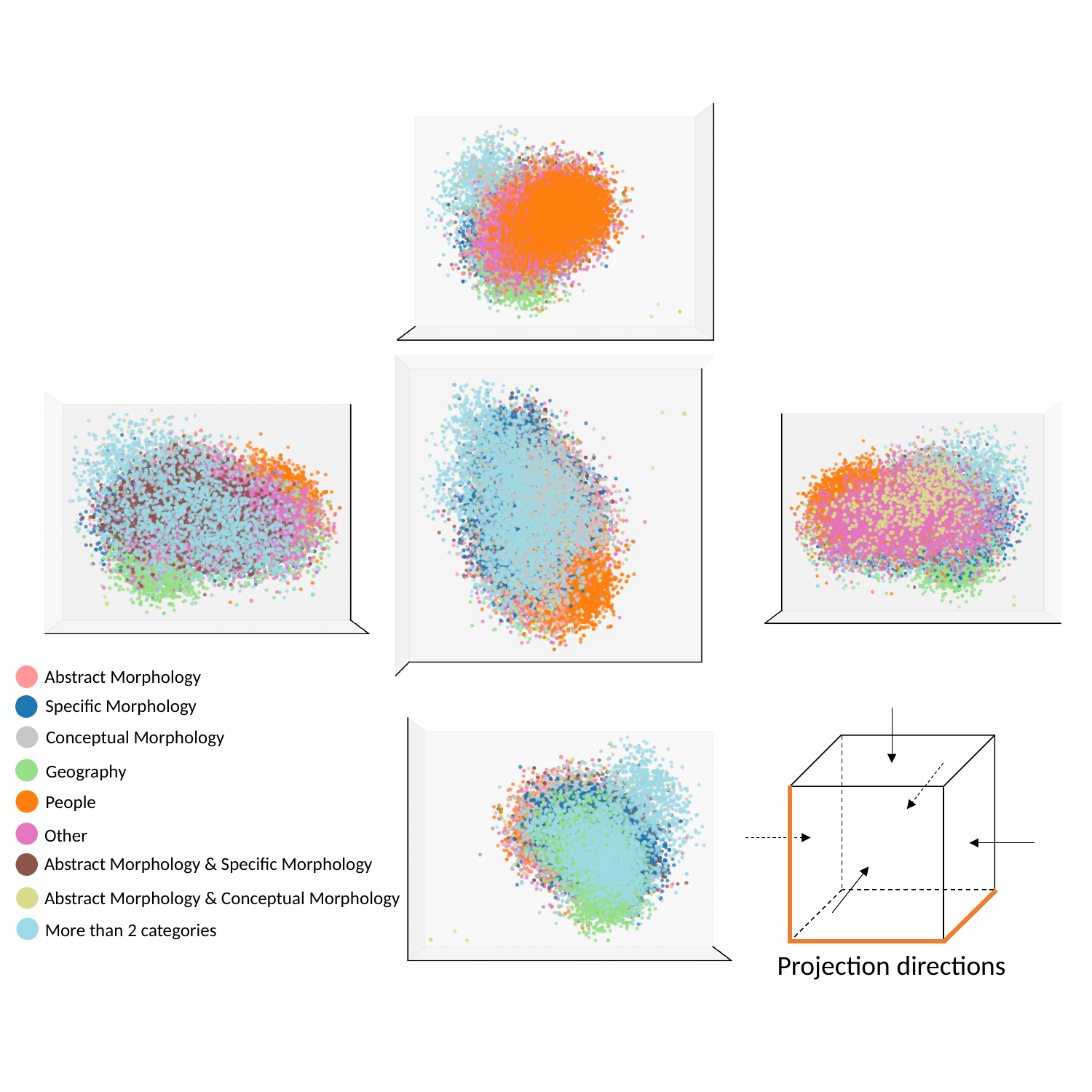

Abstract Morphology
Specific Morphology
Conceptual Morphology
Geography
People
Other
Abstract Morphology & Specific Morphology
Abstract Morphology & Conceptual Morphology
More than 2 categories
Projection directions

### Slide 4
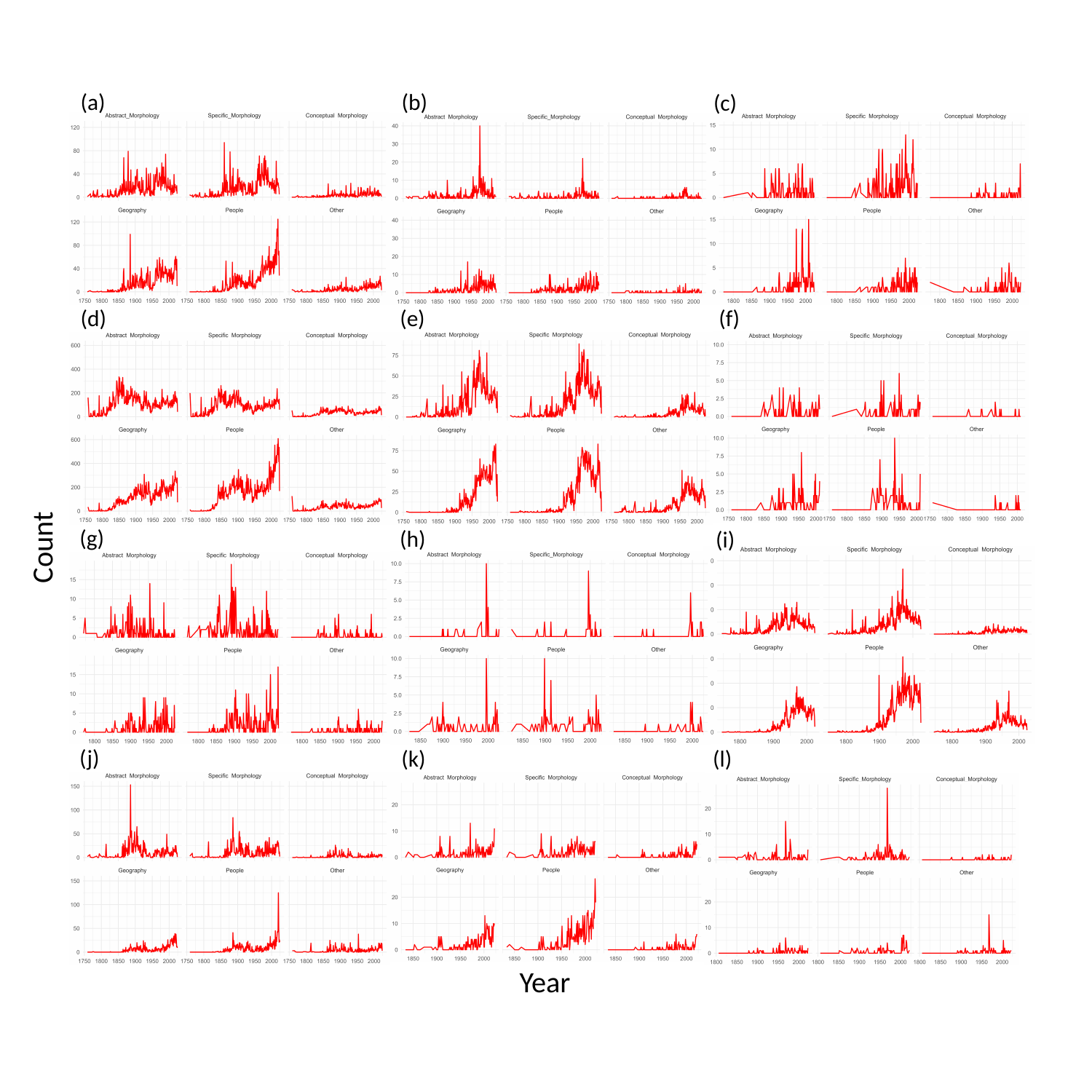

(b)
(a)
(c)
(f)
(e)
(d)
(g)
(h)
(i)
Count
(j)
(k)
(l)
Year

### Slide 5
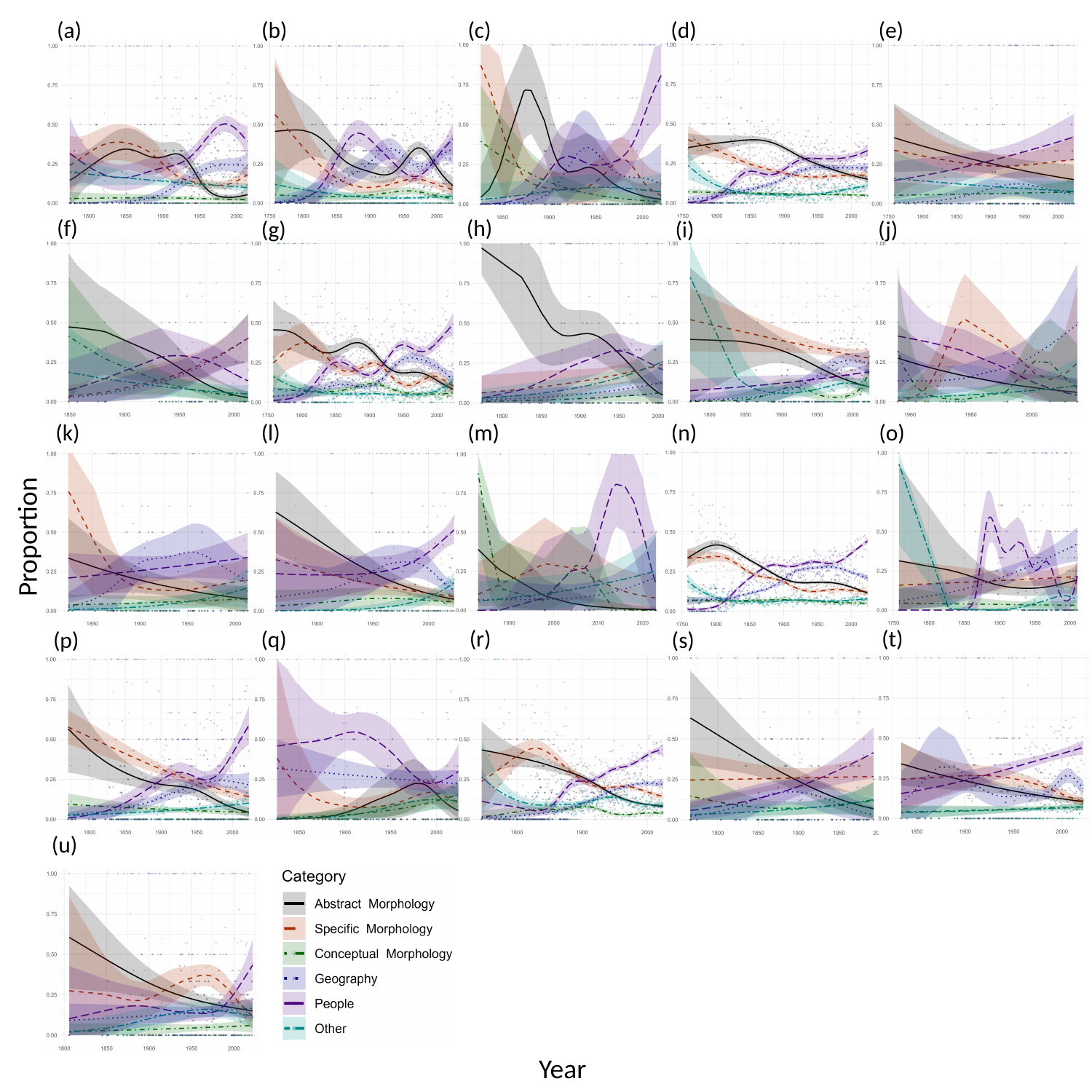

(a)
(b)
(c)
(d)
(e)
(f)
(h)
(g)
(i)
(j)
(m)
(n)
(o)
(l)
(k)
Proportion
(r)
(t)
(q)
(s)
(p)
(u)
Year
